# CropSurveyor: a scalable open-source experiment management system for distributed plant phenotyping and IoT-based crop management

**DOI:** 10.1101/451120

**Authors:** Daniel Reynolds, Joshua Ball, Alan Bauer, Simon Griffiths, Ji Zhou

## Abstract

**Background:** High-quality plant phenotyping and climate data lay the foundation of phenotypic analysis as well as genotype-by-environment interactions, which is important biological evidence not only to understand the dynamics between crop performance, genotypes, and environmental factors, but also for agronomists and farmers to monitor crops in fluctuating agricultural conditions. With the rise of Internet of Things technologies in recent years, many IoT-based remote sensing devices have been applied to phenotyping and crop monitoring that generate big plant-environment datasets every day; however, it is still technically challenging to calibrate, annotate, and aggregate big data effectively, especially when they were generated in multiple locations, and often at different scales.

**Findings:** CropSurveyor is a PHP and SQL based server platform, which provides automated data collation, storage, device and experiment management through IoT-based sensors and distributed plant phenotyping workstations. It provides a two-component solution for monitoring biological experiments and networked devices, with interfaces specifically designed for distributed IoT devices and centralised data servers. Data transfer is performed automatically though an HTTP accessible RESTful API installed on both device-side and server-side of the CropSurveyor system, which synchronise daily representative crop growth images for quick and visual-based crop assessment, as well as detailed microclimate readings for GxE studies. CropSurveyor also supports the comparison of historical and ongoing crop performance whilst different experiments are being conducted.

**Conclusions:** As an open-source experiment and data management system, CropSurveyor can be used to maintain and collate important crop performance and microclimate datasets captured by IoT sensors and distributed phenotyping installations. It provides near real-time environmental and crop growth monitoring in addition to historical and current data comparison through a single cloud-ready server system. Accessible both locally in the field through smart devices and remotely in an office using a PC, CropSurveyor has been used in wheat field experiments for prebreeding since 2016 and has the potential to enable scalable crop management and IoT-style agricultural practices in the near future.

## Background

Automated phenotyping technology has the potential to enable continuous and precise measurement of phenotypes that are key to today’s crop research [1,2]. Quantitative phenotypic traits collected through crop development are not only important evidence for biologists to understand the dynamics between crop performance, genotypes, and environmental factors (e.g. genotype-by-environment interactions, GxE), but critical for agronomists and farmers to monitor crops in fluctuating agricultural conditions [3–5]. High quality phenotyping and climate datasets lay the foundation for meaningful phenotypic analysis, which is likely to produce an accurate delineation of the genotype-to-phenotype pathway for assessing yield potential and environmental adaptation [6,7]. Presently, although many automated phenotyping platforms are capable of accumulating big plant-environment data [8], it is still technically challenging to collect, calibrate, annotate, and aggregate the data effectively, for biological experiments carried out in multiple locations, and often at different scales [9,10].

With the rise of Internet of Things (IoT) technologies and their applications in plant phenotyping [11], a number of commercial data management solutions have been developed on the base of customised hardware and proprietary software. For example, the Field Scanalyzer system (LemnaTec) employs a simple HTTP server with an SQLite database to facilitate crop monitoring and deep field phenotyping using LemnaControl and LemnaBase software [12]; Integrated Analysis Platform (LemnaTec) [13] provides an automated pipeline to combine raw image collection and metadata association for indoor phenotyping; the FieldScan system (Phenospex) [14] uses infield WiFi network to connect PlantEye^TM^ 3D laser scanners, climate sensors, and a gantry system with a PostgreSQL database to realise the scanner-to-plant phenotyping; and, PlantScreen^TM^ system (Photon Systems Instruments, PSI) manages fluorescence images and trait scores through dedicated networks and databases [15]. However, all these commercial systems require ongoing licensing maintenance and additional costs for developing new functions. It is therefore challenging for a broader plant research community to adopt and extend them easily in order to meet the growing needs of today’s crop research [10].

Recently, some research-based systems have also been introduced to the scientific community. For example, PhotosynQ software manages data collection and storage through a handheld device called MultispeQ [16]. It uses Bluetooth to retrieve leaf surface images, environmental and geolocational data collected by MultispeQ and stores them in a mobile phone or a laptop. The system requires manual interference for data synchronisation and centralised analysis via onsite workstations or cloud-based servers. Hence, it is tailored for small-scale and qualitative phenotyping tasks. BreedVision is another system that gathers data through a network-based HTTP server [17]. Mounting multiple sensors on a tractor, BreedVision is used to carry out field phenotyping for wheat breeding. Sensors communicate to a SQL database running in an embedded system. However, this platform is designed for bespoke hardware and does not provide an open application programming interface (API), indicating that it is incompatible with external hardware and software. Solely for collecting climate datasets, the PANGEA architecture [18] was successfully established to network large numbers of connections (e.g. wireless sensor networks, WSN) for agricultural practises [19]. This system has been used to integrate large-scale WSN installations through open and distributed smart device interfaces. However, it cannot handle image-based datasets and thus limits its applications in phenomics driven crop research. Lately, a comprehensive and open-source Phenotyping Hybrid Information System (PHIS) has been developed by INRA [20]. PHIS system aims to provide a platform to enable data tracing and reanalysis of phenomic data collected on thousands of plants, sensors and events. It can identify and retrieve objects, traits and relations via ontologies and semantics. Because the PHIS system needs to incorporate many external phenotyping and modelling systems, it is heavyweight and mainly focuses on post-experimental data integration and analysis.

The above industrial and academic efforts identify the need to develop a scalable and openly available data management system. It needs to handle different types of datasets acquired in automated plant phenotyping experiments. To integrate data transfer, calibration, annotation and aggregation effectively, such a system should be flexible for changeable experimental designs and expandable with third-party hardware and external software. More importantly, the system needs to enable users to closely monitor experiments conducted in different locations whilst the experiments are being conducted.

With these design requirements in mind, we developed CropSurveyor, an IoT-based data management system that is easy to use and flexible to deploy in diverse experimental scenarios. CropSurveyor is a scalable and open-source software system, which provides diverse interfacing options for the community to adopt and extend. We followed a distributed IoT systems design during the development, so that experimental, phenotypic, and environmental data collected from infield and indoor experiments could be integrated efficiently. The system provides a unified web interface for users to oversee data collection, calibration and storage on a regular basis. Through our three-year wheat prebreeding experiments (2016-2018) [21], a powerful visualisation component and a flexible data/experiment management solution has been established. Equipped with CropSurveyor, users can now closely monitor different experiments, ongoing and historic, running in different locations. Furthermore, the modulated software architecture has made it possible to change scale and performance for new experimental needs. To our knowledge, the research-based CropSurveyor system has the potential to significantly contribute towards dynamic data collection and experimental management, for both plant phenotyping and crop GxE studies.

## Findings

IoT is a fast-growing field. IoT-based sensors are generating terabytes of data for crop research and agriculture services everyday [22]. Because the existing data management solutions heavily rely on bespoke data collection approaches, they cannot be easily adopted and extended. Also, most of the solutions require the construction of a centralised management system, which would not resolve the problem of scalability and accessibility, because the distributed nature of IoT technologies and the centralised data administration infrastructure are likely to confound each other. Instead, we developed a two-component solution. The first part of this is a device-side system that is lightweight and capable of interacting directly with distributed IoT devices, ensuring onboard data standardisation and data collection. The second component is a server-side system that collates and stores image‐ and sensor-based datasets, with SQL as the back-end. This server-side system is comprehensive and responsible for visualising dynamic crop-environment data collected during experiments. Combining both parts, the open-source CropSurveyor system is capable of bringing scalability and flexibility to users.

### The systems design

The two-component systems design of CropSurveyor is shown in Fig. 1. We used a Python-based web framework, Flask [23,24], as the base for the device-side services. The main reason for this choice is that Python, a high-level programming language widely used by the scientific community, can interact with many single-board computers (e.g. a *Raspberry Pi* computer) commonly embedded in distributed IoT sensors and/or phenotyping devices. This framework administers onboard data storage and establishes a lightweight server for web-based interactions (Fig. 1A). As Flask is hardware independent, the approach can be applied to any hardware that supports Python. Additional services such as Linux *crontab* scheduling system, dynamic host configuration protocol (DHCP, used for establishing self-operating WiFi network), and virtual network computing (VNC) services can be easily added or removed to maintain the simplicity of the device-side system.

**Figure 1:**
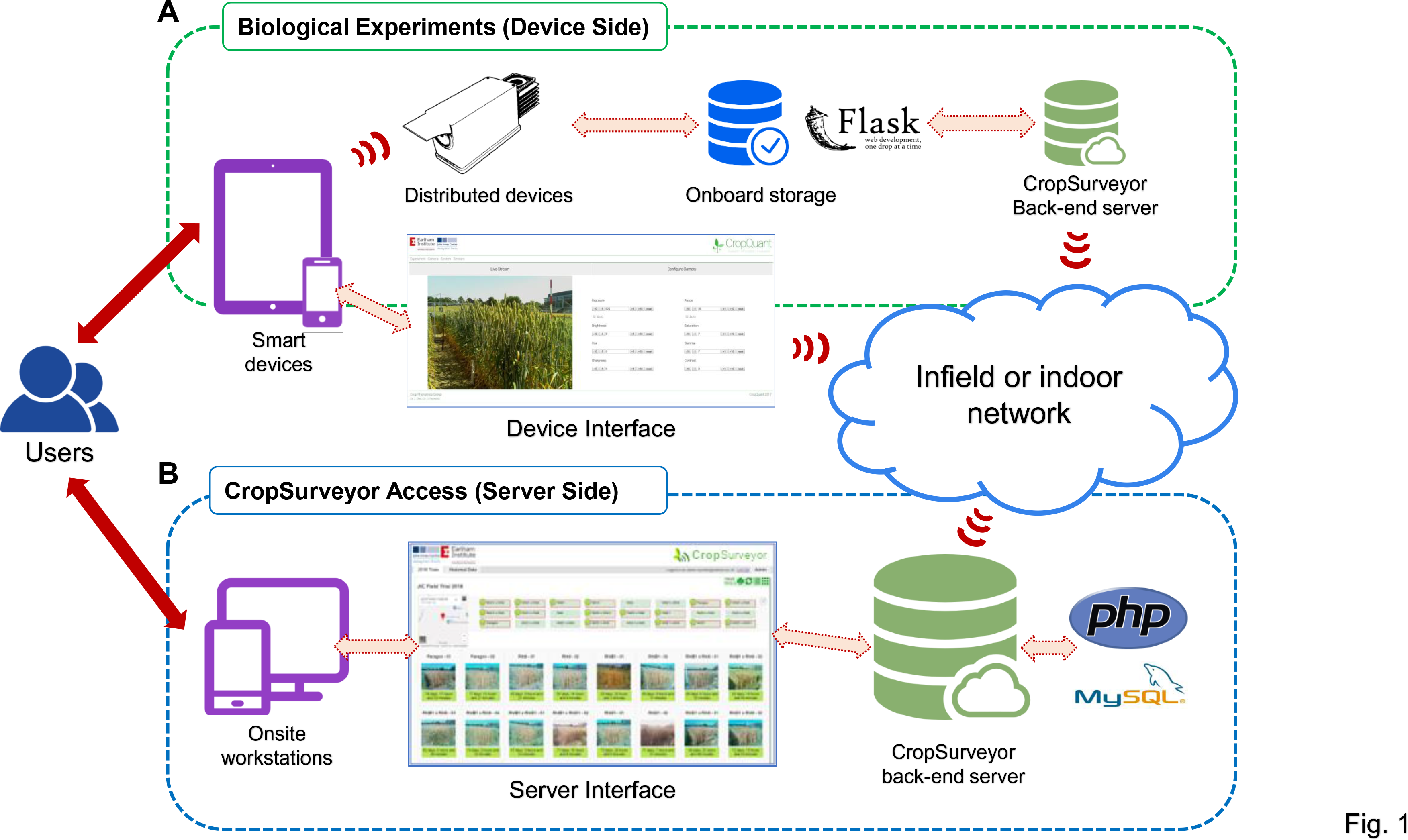
A deployment diagram of the CropSurveyor system in biological experiments. **(A)** CropSurveyor facilitates users to interact with distributed infield or indoor phenotyping workstations using wired (e.g. ethernet cables) or wireless connection (e.g. smart devices’ WiFi). The CropSurveyor client running on distributed workstations supports remote systems interactions and onboard data management. **(B)** Users can connect, monitor and administer experiments using CropSurveyor server in real time. Through dedicated networks, the CropSurveyor back-end server collates and integrates large image‐ and sensor-based phenotyping datasets in an SQL database.

Powered by PHP5+ [25] and MySQL [26], the device-side system can facilitate real-time interactions between smart devices (e.g. smartphones and tablets) and IoT devices. The graphic user interface (GUI) was developed using PHP and JavaScript, which can be opened in a web browser such as Chrome and Firefox on any smart device. A PHP-based RESTful API [27] was adopted to regulate hourly client-server communications. A lightweight SQL server, MariaDB [28], was used for collecting and storing different formats of datasets, including images, climate sensors, and experimental settings. The device-side system can also be used to initiate a live video streaming for users to deploy infield or indoor phenotyping devices (Supplementary Fig. 1), so that an experiment can be initiated or terminated via a smartphone or a tablet. Also, the GUI allows users to enter metadata including trials, experiments (e.g. genotypes, treatments and biological replicates), and brief description, while phenotyping devices are being installed. The distributed IoT-based design has massively improved the mobility and flexibility for conducting phenotyping tasks.

The server-side system bridges the connection between data aggregation and cloud-based interfacing (Fig. 1B). This approach facilitates biological datasets acquired at different locations to be synchronised with a centralised server for detailed traits analyses and decision making in crop management. PHP5+ was used to develop the system that supports Apache and an SQL server such as MySQL [26]. The server-side system initiates regular updates of the status of each distributed IoT device with information such as online or offline status of the device, operational mode, representative daily images, micro-climate readings, and the usage of computing resources (i.e. CPU and memory). Since 2017, the two-component CropSurveyor system has been successfully applied to monitor indoor wheat speed breeding [29] and infield wheat prebreeding simultaneously (Supplementary Fig. 2).

### An MVC architecture

Whilst CropSurveyor is designed to allow users with no technical background to use, the installation of the system still requires an IT technician to complete (see Additional File 1). To install the system, a functioning PHP and SQL server is required. Also, as it runs on a network-enabled web server, a network infrastructure is required to properly function CropSurveyor (Fig. 2). However, due to the rural location of many crop research experiments, it is often expensive and unfeasible to install wired or wireless networks in some experimental sites. Hence, our solution is to establish an ad-hoc and self-managed network through USB WiFi dongles mounted on IoT devices, e.g. a distributed CropQuant phenotyping workstation [21], so CropSurveyor can transfer data between distributed IoT devices and a central server. The self-managed network can be either a Star or a Mesh network topology, enabling peer-to-peer HTTP accessing points to network IoT devices for data calibration and synchronisation in the field (Fig. 2A), or to establish a direct link between a smart device and a phenotyping workstation (Fig. 2B). After correlating and collecting all data from the device side, the system will then transfer the data to the server-side powered by a central server, where users could oversee different experiments at near real-time (Fig. 2C).

**Figure 2:**
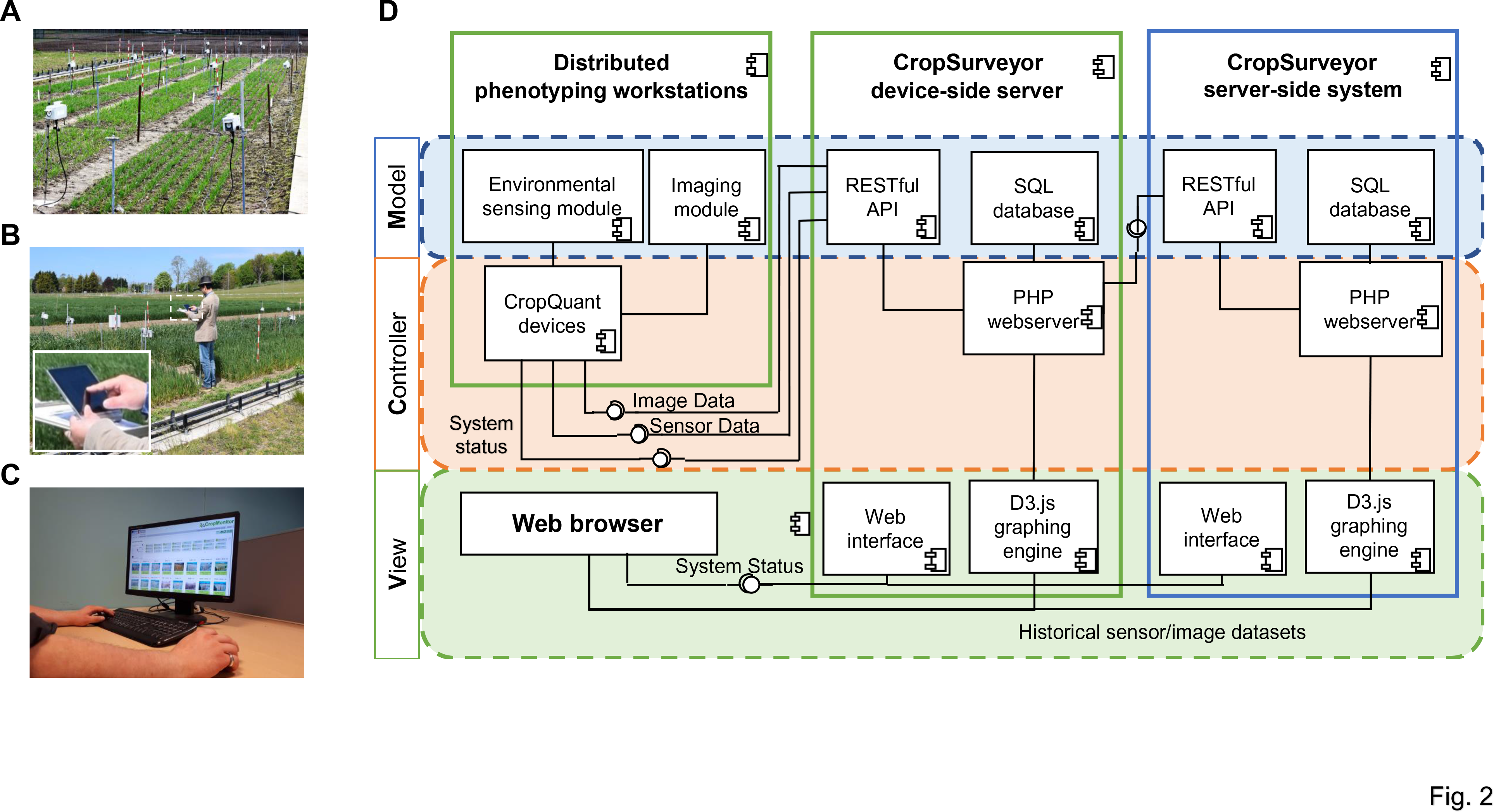
A component diagram of the real-world deployment and application of the CropSurveyor system. **(A)** IoT phenotyping workstations installed at Norwich Research Park. Distributed nodes are connected by the cloud-based CropSurveyor system. **(B)** Infield phenotyping devices can be directly accessed and controlled through the local CropSurveyor client directly in the field by using a smart device. **(C)** CropSurveyor can be used remotely to manage ongoing experiments through an accessible web interface. **(D)** A detailed component diagram showing the MVC design of the CropSurveyor system and the interface between infield/indoor phenotyping workstations, local CropSurveyor server, cloud-based server and user interactions. The data input is through a RESTful API, responsible for transferring data between servers and enabling interactions through a web-based user interface.

When implementing the CropSurveyor system, we followed Model-view-controller (MVC) software architecture, dividing the system into three interconnected parts to separate internal information flows based on how they are presented to the user [30]. Using the MVC pattern to interface different parts of the CropSurveyor system, not only source code of both device-side and server-side systems can be reused, we could also enable modulated parallel software development, while biological experiments were still ongoing (Fig. 2D).

To enable data standardisation and integration, a RESTful API was implemented that accepts image‐ and sensor-based datasets and IoT device status updates in JSON format. All interactions between devices and the server are authenticated using a pre-shared key pair to ensure that datasets collected are from a trusted source. The RESTful design strategy ensures that all data requested for transaction is contained within a single request, allowing devices to compile all information into one JSON object and then transmit through an HTTP POST request. The *Model* implementation allows us to determine dynamic data structure, as well as how to manage logic and rules of the CropSurveyor system. The entity–relationship model (ER diagram) used for establishing the database including entity types and specifies relationships between the entity types can be seen in Supplementary Fig. 3.

Based on PHP server (Apache tested) and SQL server (MySQL and MariaDB tested), the *Controller* component responds to user input and internal interactions on the data model. The controller receives image, sensor and system status as the input data flows, validates them, and then passes them to the model component, first on distributed device-side server and then transmitted to a globally accessible server-side server, which mirrors the input data. Internet connections are required, if the input datasets need to be transferred from a field experiment site to onsite servers. The form of data transmission can be either wired ethernet or Wi-Fi network. The Controller administers data collation between device-side and server-side by mimicking the device API call to the higher-level server API, at the time of device request is programmed.

The *View* component presents the data model and user interactions in two formats. First, through an active HTTP connection and D3.js graphing engine [30], users can access distributed IoT devices via web browsers (Chrome and Firefox tested) installed on any smart device, in the field or in greenhouses. The device-side CropSurveyor provides a tailored GUI interface, within which users can deploy (see Additional File 1), monitor, assess and download captured data on demand. Second, the device-side system synchronises with the server at regular intervals, based on which CropSurveyor provides a more comprehensive GUI to present both experimental and technical status of ongoing experiments. The device-side system is designed to be distributed. So, if a given IoT device cannot make a direct internet connection for any reasons, the device-side system will enable local data storage as a server node. After the networking is re-established, the system can then forward collected data automatically (the onboard USB memory stick normally can store 30 days’ image and sensor data).

### Experiment and data management

Monitoring dyanmic plant phenotypes such as height, growth rate, growth stages, and associated climate conditions in biological experiments can be a laborious and time-consuming task. It is even more challenging if we need to calibrate and verify datasets collected via devices deployed in different sites. In particular, low-quality and missing data often leads to analysis errors and unusable results, normally identified after the completion of a given experiment [31]. Hence, the server-side CropSurveyor system was designed to oversee ongoing experiments based on representative daily images, hourly sensor data collected from each phenotyping device, as well as experimental settings such as genotype, treatment, drilling date, plot position and biological replicate.

The interfaces of experiment and data management are presented in Fig. 3, which integrate experiment location, plot map, and crop/experiment/device information to enable quick cross-referencing and facilitate management decisions during the experiment. As shown in Fig. 3A, for a given experiment, the grid view of the server-side system provides a set of device nodes showing GPS-tagged project geolocation, identifiers of installed phenotyping devices, representative daily images of monitored plots, and colour coded status indicator displaying the operation mode of each device. CropSurveyor reads the device-side server’s GPS coordinates and presents the geolocation in an embedded Google Map for users to locate the experiment. In addition to the GPS location of the experiment, an embedded plot map is also provided demonstrating individual device position in the field or in greenhouses together with colour coded status markers on the relevant plots to quickly indicate whether extra attention is needed (e.g. green for operating, amber for idle, and red for device termination or operational error). These markers in the plot map can be clicked, which will bring the user to the detailed view of individual device (Fig. 4). Each phenotyping device uploads a daily representative image of the monitored plant or plot. The resolution of the image is 640×480 pixels, downsized from 2592×1944 pixels to enable constant data transmission for large-scale device-server data synchronisation. The image is automatically selected based on file size, intensity, and image clarity. The grid view of these representative image is used as a snapshot of the experiment, so that users can quickly assess plant growth and performance of each genotype without regularly walking in the field during the growing season.

**Figure 3:**
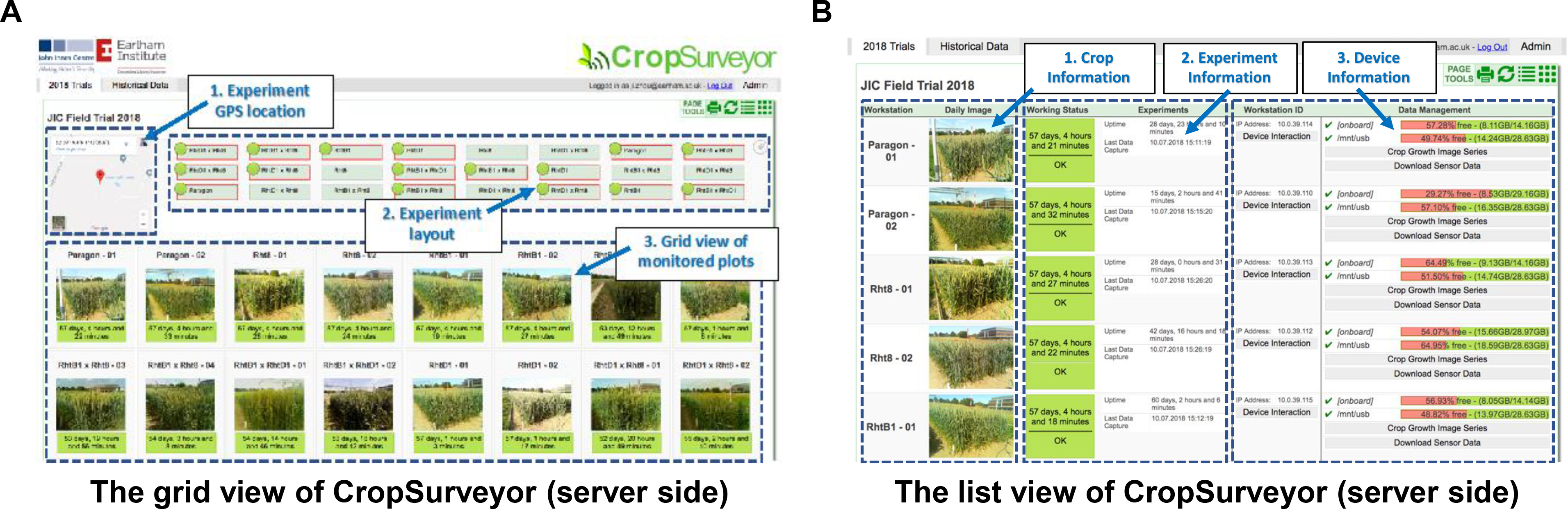
System views of CropSurveyor’s user interface. **(A)** The user interface is accessible through a web browser on any computing device. The grid view of the system is designed for integrating key experimental information, showing geolocation of field experiments, experiment layout, monitored plots and genotypes, experiment duration, and representative daily images of the monitored genotypes. **(B)** The list view shows detailed statistics of all monitored crops in a given experiment, including crop information (genotypes and representative images), experimental information, and workstation information such as workstation ID, its storage, IP address, image and sensor data download, and device interaction function devices flask-based HTTP interface. This view is more useful from a system management perspective.

The list view provides a table of status that incorporates crop information with experiment and device details (Fig. 3B). This view is mainly used for project maintenance proposes, which contains three sections. First, similar to the grid view, crop information identifier lists phenotyping devices installed in the experiment. Second, experiment information includes a coloured status indicator to display the operational mode of a given device, the experiment duration of a given device, and the latest timestamp of data synchronisation. Device uptime (i.e. experiment duration) is computed using the device’s internal clock (the Linux uptime command) and the time when the latest image is captured. Third, device information shows: (1) each device’s onboard storage, using filled bars to indicate the percentage of space left in gigabytes (GB) based on regular 30-minute updates; (2) buttons to download image‐ (“Crop Growth Image Series”, in monthly Zip archives) and sensor-based (“Download Sensor Data”, in a CSV file) datasets collated during the experiment from the SQL database; and (3) device interaction buttons, providing direct device control and configuration via Secure Shell (SSH) or VNC.

### Continuous microclimate visualisation

Microclimate is an important evidence for crop scientist to monitor radiation/ambient/soil variation in different locations over the whole experiment site, which closely connects with the performance at both plant and plot levels [32]. To facilitate the monitoring of microclimate during the experiment, a comprehensive visualisation function has been developed (Fig. 4). By accessing an individual device’s detail page, collected environmental factors can be viewed as individual line charts along with the device information. IoT-based climate sensor readings are logged with the central server and then indexed by device and location, allowing near real-time microclimate readings (30-minute updates) of monitored regions. The visualisation is done in the web browser using the D3 JavaScript library. In our case, we can soundly retrieve readings such as device temperature (to assess device performance), ambient relative humidity, ambient temperature (Fig. 4A), light levels (based on light intensity), soil temperature and moisture (Fig. 4B). The microclimate datasets acquired from multiple locations across the field can also be used for data calibration to generate a normalised and highly reliable environmental reading of the experimental site.

**Figure 4:**
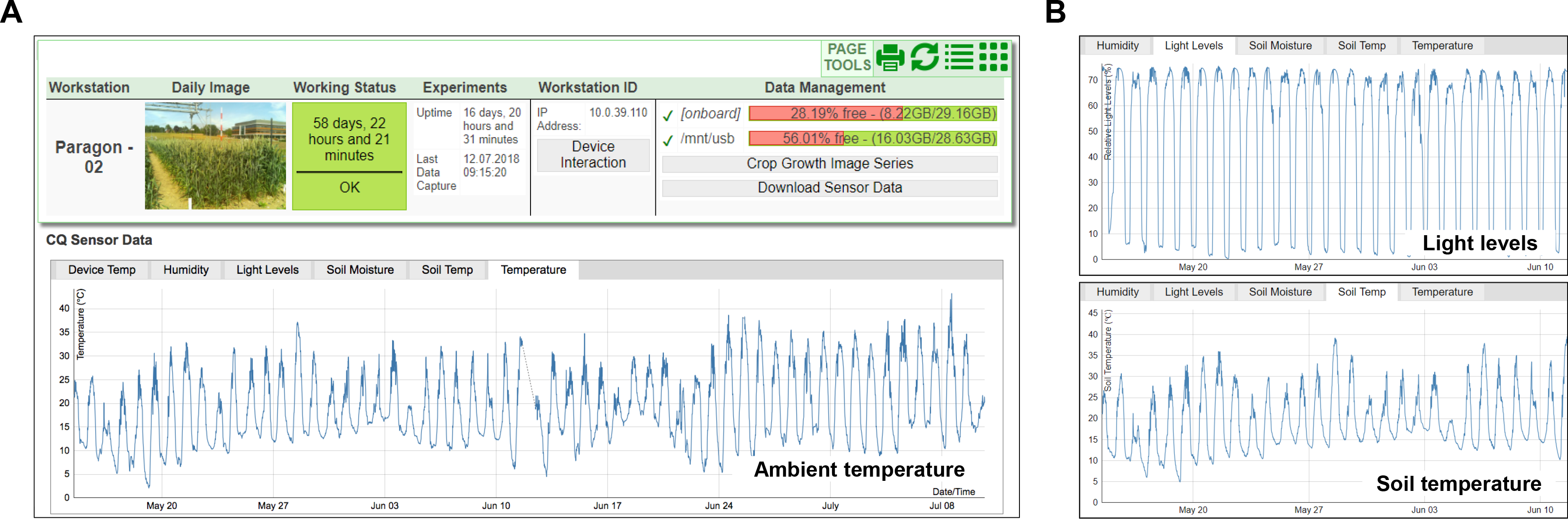
An individual view of a given genotype monitored by the CropSurveyor system. **(A)** The individual view of a monitored genotype/plot accessible through the CropSurveyor user interface, detailing device and experiment information together with captured environmental sensor data. **(B)** Web-based graph visualisation of hourly sensor readings during a given experiment, showing ambient temperature, ambient humidity, field lighting, soil moisture, and soil temperature variation in the plot region.

### Applications in wheat field experiments

A key element of modern agriculture is to closely monitor dynamic crop performance and agricultural conditions to predict and plan crop production [33]. Plant breeding and GxE studies also rely on high-quality and high-frequency crop-environment data to produce accurate growth models for yield and quality prediction [34,35]. Following this approach, CropSurveyor provides users with quick access to all environmental factors recorded by each distributed phenotyping device during the growing season. Together with the position of a given device, seasonal microclimate datasets can form a dynamic growth condition map showing environmental conditions and variance in a given field (Fig. 5).

**Figure 5:**
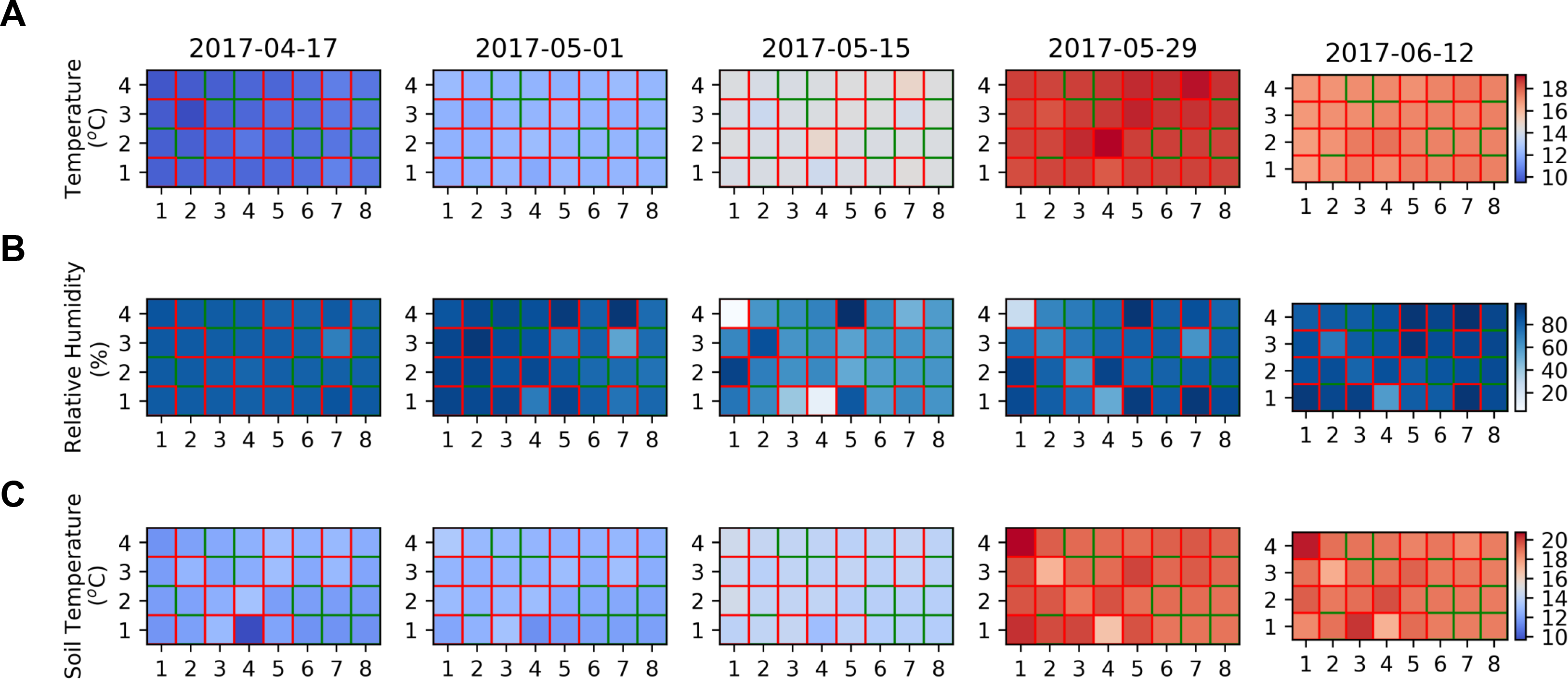
The infield microclimate conditions collated by the CropSurveyor system. **(A, B)** A Heat map of ambient sensor reading of temperature and relative humidity recorded during the growing season. Each cell represents an individual plot in the 2017 field experiment. Real sensor reading outlined in red and interpolated values outlined in green. **(C)** A Heat map of soil-based sensor reading of soil temperature recorded during the growing season.

In a 253-day field experiment of 32 wheat genotypes within the single genetic background of Paragon (a UK spring wheat variety) accomplished in 2017, we have installed 16 CropQuant field phenotyping workstations to monitor six-metre wheat plots to collect continuous crop growth image series as well as associated microclimate conditions such as ambient temperature, relative humidity, light levels, soil temperature and soil humidity. When the environmental data was being collated, a field map of dynamic microclimate conditions at key growth stages (i.e. from early booting to early grain filling, 56 days) was gradually produced, showing the increase in ambient temperature (Fig. 5A), the variation of ambient moisture levels (Fig. 5B), and the steady increase of soil temperature (Fig. 5C), during the 56-day period. To simplify the presentation, the microclimate heatmap was presented with data at 14-day intervals, where wheat plots installed with IoT sensors were outlined with red colour and plots without sensors were outlined with green colour, where climate data was produced through data interpolation methods based on adjacent readings (Fig. 5). The period of the interval can be flexibly changed, and the microclimate readings are retrievable as soon as data synchronisation is finished (see Supplementary Fig.4 for daily data presentation). Furthermore, the climate datasets can be used for cross-validating the soundness of infield IoT sensors, for example, whether soil temperature correlates with ambient temperature (Supplementary Fig. 4A); and why readings from distributed low-cost sensors could provide more representative information of the field in comparison with an expensive central weather station in the field (Supplementary Fig. 4B).

Utilising this approach, dynamic environmental conditions throughout a field can be recorded with very low-cost climate sensors, which can then be scaled up through interpolation methods to cover regions without sensors. Through wheat field experiments between 2016 and 2018 at Norwich Research Park, we believe that distributed IoT sensors together with the CropSurveyor system are capable of providing invaluable crop monitoring and management data in near real-time.

### Comparison between multi-year experiments

CropSurveyor not only provides tools for monitoring ongoing infield and indoor experiments, but also supplies toolkits to reference and download historical datasets. An important function in crop research is the ability to compare collected results with past experiments. To this end CropSurveyor stores all image and sensor data and manages these historical datasets with easy reference and access (Fig. 6). Historical datasets can be retrieved through the frontpage similar to ongoing experiments (multiple projects can be administered by CropSurveyor simultaneously). After opening a completed project, users can display the GPS-tagged geolocation of the project and devices used in the project together with project references (Fig. 6A). By clicking a specific plot within the experimental field, CropSurveyor can directly reference environmental and image datasets in the plot, a view with device name, date of last capture and last image taken by the IoT device (Fig. 6B). If users want to revisit previous datasets in the project, they can download both sensor data packages and/or growth image series in monthly archives by clicking the archive links (Fig. 6C). This design enables a unified platform to facilitate both ongoing and historical data management to assist in-experiment and post-experiment data analysis.

**Figure 6:**
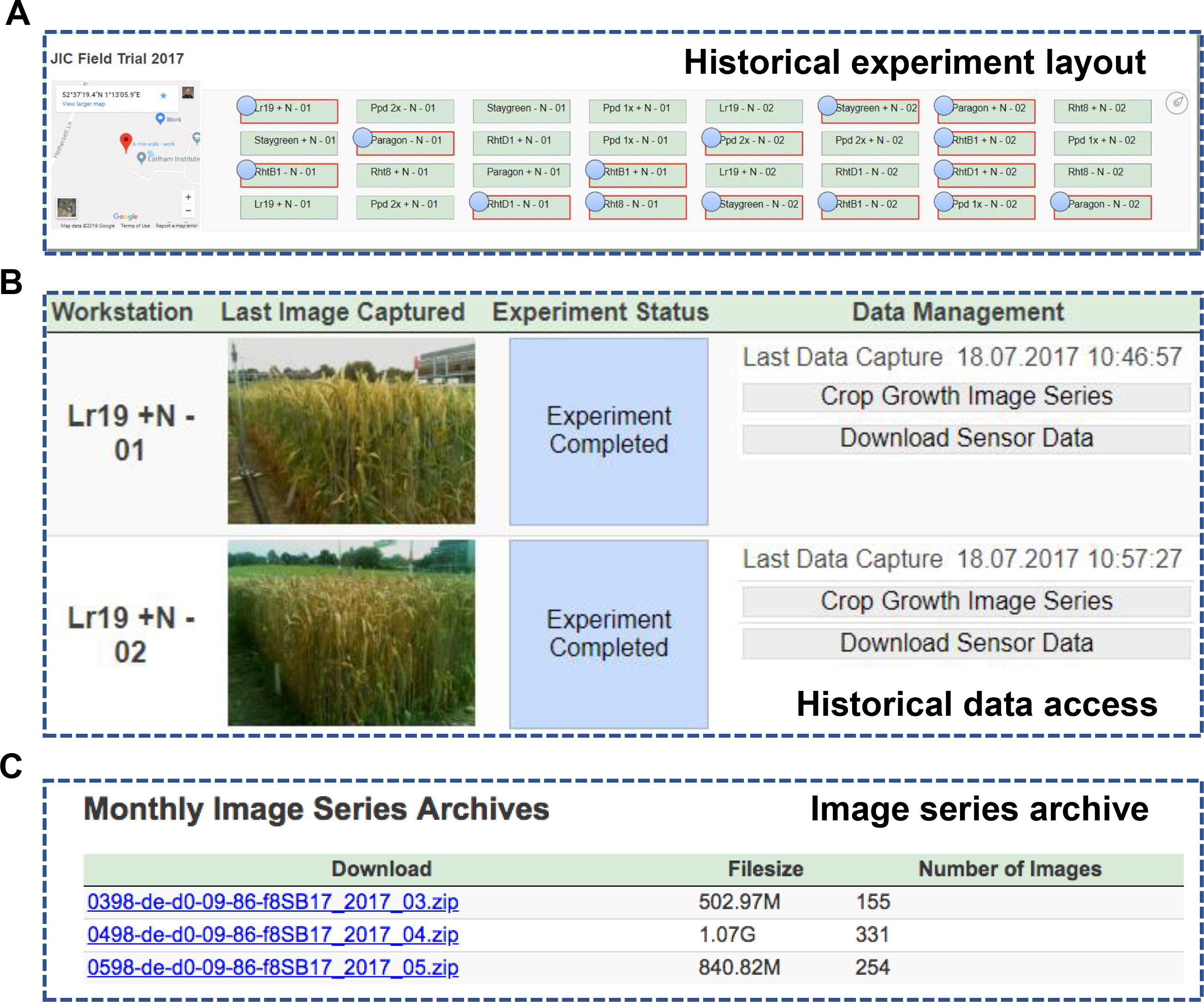
Historical experiment data access. **(A)** The CropSurveyor system provides access to historical experimental datasets, including the geolocation of a given project and genotypes/plots monitored in the completed project. **(B)** In a completed project, the last image captured in the experiment as well as historical image‐ and sensor datasets can be downloaded. **(C)** The download links for monthly image series archived in cloud.

## Discussion and outlook

The continuing challenge of global food security caused by fluctuating environments and a narrower range of genetic variation of modern crops requires innovative thoughts and technologies to improve crop productivity and sustainability [2,36,37]. As European infrastructures for sustainable agriculture (e.g. EMPHASIS and AnaEE) have identified, openly shareable solutions built on widely accessible digital infrastructures are likely to provide an effective solution to address the challenge by integrating novel scientific concepts, sensors and models [38,39]. The IoT-based CropSurveyor system presented here is scalable and open-source, providing the scientific community various interfacing options to adopt and extend. The openly available platform integrates the archiving and collation of high-frequency environmental data and crop images automatically, which can be used for both phenotypic analyses as well as agricultural decision making. By associating environmental conditions directly with crop growth data, we trust that the system is capable of forming a sound base for reliable GxE studies. More importantly, CropSurveyor provides geolocation and remote sensor readings of current and historical experiments, a comprehensive solution to enable multi-site and multi-year cross-referencing of traits analyses as well as crop performance monitoring.

Because CropSurveyor facilitates the real-time distributed access of microclimate conditions and crop imagery (through live video streaming) on-demand in the field or in greenhouses, either through a smart device or an office PC, users can make a quick decision of crop performance, growth stages, and plot conditions of any monitored locations in a given experiment, field, or site. More importantly, automatic data transmission allows a centralised data and experiment management, which means that the system can be scaled up to the national scale if a broader IoT in agriculture infrastructure is in place. As collected data is annotated and pre-selected on distributed phenotyping or IoT-based devices, only standardised crop-environment datasets are collected from different experiments to support detailed analysis and meaningful cross-referencing. Furthermore, openly sharing results from different sites and different experiments will enable crop researchers, breeders, and farmers to gain great benefits, for example, predicting and prewarning disease spread at the national scale so that early adoption of preventative measures can be arranged.

Presently, many governments are shifting their focuses towards innovative technologies to modernise crop and agricultural research. The UK Government, for example, has invested heavily in IoT-based technologies to address challenges on yield production, food traceability, environmental challenges, incompatibility, and lack of infrastructure [40]. We believe that CropSurveyor can also address some of the current challenges directly. For example, by logging historical data and annotating crop growth and environmental effects within monitored fields can increase crop traceability. To reduce the overall use of agrochemicals as part of a precision farming strategy [41,42], CropSurveyor can be used to identify the appropriate timing and areas for chemical application together with infield imaging and ambient sensors. Water is in limited supply for large regions of the globe and the reduction of unnecessary irrigation would be of large benefit to the cost-effectiveness of agriculture [43,44]. As discussed previously, CropSurveyor is built in with near real-time environment monitoring mechanisms including soil temperature, soil moisture levels, and ambient humidity. Hence, it provides information crucial to make decisions and targeting irrigation in timing and location. Additionally, by linking extra climate sensors with IoT devices, further environmental readings can be extended in CropSurveyor for growing agricultural needs.

Besides the near real-time environmental and crop growth monitoring, historic and current datasets collated in a central system can also deliver predictive powers. An example of potentially predictable situations is the “Smith Period” for predicting Late Blight in potato crops [45]. Late Blight is shown to be more likely to occur during a “Smith Period”, which is defined by a period of two or more days with a minimum temperature of 10°C and a humidity of 90%, or above for at least 11 hours in each day. Having direct access to dynamic sensor readings on the CropSurveyor can allow the monitoring of specific environmental patterns much easier and thus establish an important tool to inform farmers and growers to apply fungicides and chemical treatments to the appropriate areas. Based on this potential development, CropSurveyor is potentially able to serve sustainable agriculture and environmentally friendliness of food production under today’s changeable climates.

### Future Development

To establish a data and experiment management system that is scalable and usable on regional, national or even global crop research and agricultural practices, we believe that CropSurveyor in connection with distributed IoT sensors can meet the future demand of usability and scalability, with some further development. One area of expansion is in scalability. The system is currently tested on local server with a direct network connection to at least one of the distributed nodes. To allow the expansion at a larger, national, or even global scale, the reliance on maintained servers would be less effective than a true cloud enabled service. Hence, by moving the CropSurveyor system to a globally accessible cloud server with cloud enabled distributed storage is a feasible approach, as the requirements for institutions and agricultural practitioners to maintain servers and storage are removed. Given the lack of network infrastructure in rural areas in many countries, the addition of 3G or 4G mobile data networks to key distributed nodes in the field can improve the infield network, upon which the data communication of a large number of Agri-Tech devices can be relied.

Another prohibitive factor in IoT in agriculture is the quantity and costs of IoT devices required to cover an entire field. Based on our three-year field experiments, we believe that installing sensors and phenotyping workstations to cover every area in the field is unnecessary. Fig. 5 shows that the data interpolation approach we have applied to generate microclimate readings between randomly positioned stations to model the effect of environmental variation in the whole experimental field.

This approach of subsampling produced high-quality environmental readings, which we believe could be key to the effective and feasible use of IoT in agricultural. Additionally, with the development of national IoT infrastructure, the similar subsampling idea can be expanded to a larger and multi-site level, which can then truly help inform decision in crop research and agricultural practices at the national level, across a country’s arable land.

## Availability and requirements

Project name: CropSurveyor for wheat prebreeding in Designing Future Wheat

Project home page: https://github.com/Crop-Phenomics-Group/cropsurveyor/releases

Operating system(s): Platform independent

Programming language: Python, PHP, JavaScript, SQL

Requirements: Apache (or other PHP5+) server, MySQL (or other SQL) server, a recent version of Chrome, Firefox, or Safari

License: BSD-3-Clause available at: https://opensource.org/licenses/BSD-3-Clause

## Availability of supporting data

The datasets supporting the results presented here is available at https://github.com/Crop-Phenomics-Group/cropsurveyor/releases. Snapshots of the code and other supporting data are also openly available in the GitHub repository.

## Additional files

Additional File 1.docx

MS Word Document (.docx)

CropSurveyor Installation Instructions and Interface Details

Additional file gives step-by-step instructions for initialising the system through an existing PHP webserver with SQL database, details of RESTful API required fields necessary for device interaction, and addition detail of distributed installation and database integration.

Additional File 2.html

Web Page (.html)

Algorithm to generate plotted figures

Additional file contains full python code to replicate plotted figures within the paper, displayed within an exported iPython notebook. All datasets shown within the plotted figures of the paper are available at the project GitHub repository.

## Abbreviations

AnaEE: Analysis and Experimentation on Ecosystems; API: Application Programming Interface; CPU: Central Processing Unit; CSV: Comma Separated value; DHCP: Dynamic Host Configuration Protocol; ER: Entity Relationship; GB: Gigabyte; GPS: Global Positioning System; GUI: Graphical User Interface; GxE: Genotype by Environment; HTTP: Hypertext Transfer Protocol; IoT: Internet of Things; IT: Information Technology; JSON: JavaScript Object Notation; MVC: Model View Controller; PHIS: Phenotyping Hybrid Information System; PHP: PHP Hypertext Pre-processor; PSI: Photon Systems Instruments; SQL: Structured Query Language; UK: United Kingdom; USB: Universal Serial Bus; VNC: Virtual Network Computing; WSN: Wireless Sensor Network

## Author contributions

JZ and DR wrote the manuscript. SG provided wheat expertise and germplasm. JZ and SG designed the experiment. DR and JZ designed the CropSurveyor system. DR developed the system. JZ, JB and AB tested and packaged the system. JZ and DR performed the data analysis. DR, JB and JZ deployed hardware and software for experiments. All authors read and approved the final manuscript.

## Acknowledgements

The authors would like to thank all members of the Zhou laboratory at EI and Nanjing Agricultural University for fruitful discussions. We thank The NBI Partnership (NBIP) computing team’s support. We also thank researchers at John Innes Centre and UEA for constructive discussions.

---

**Supplementary Figure 1:**
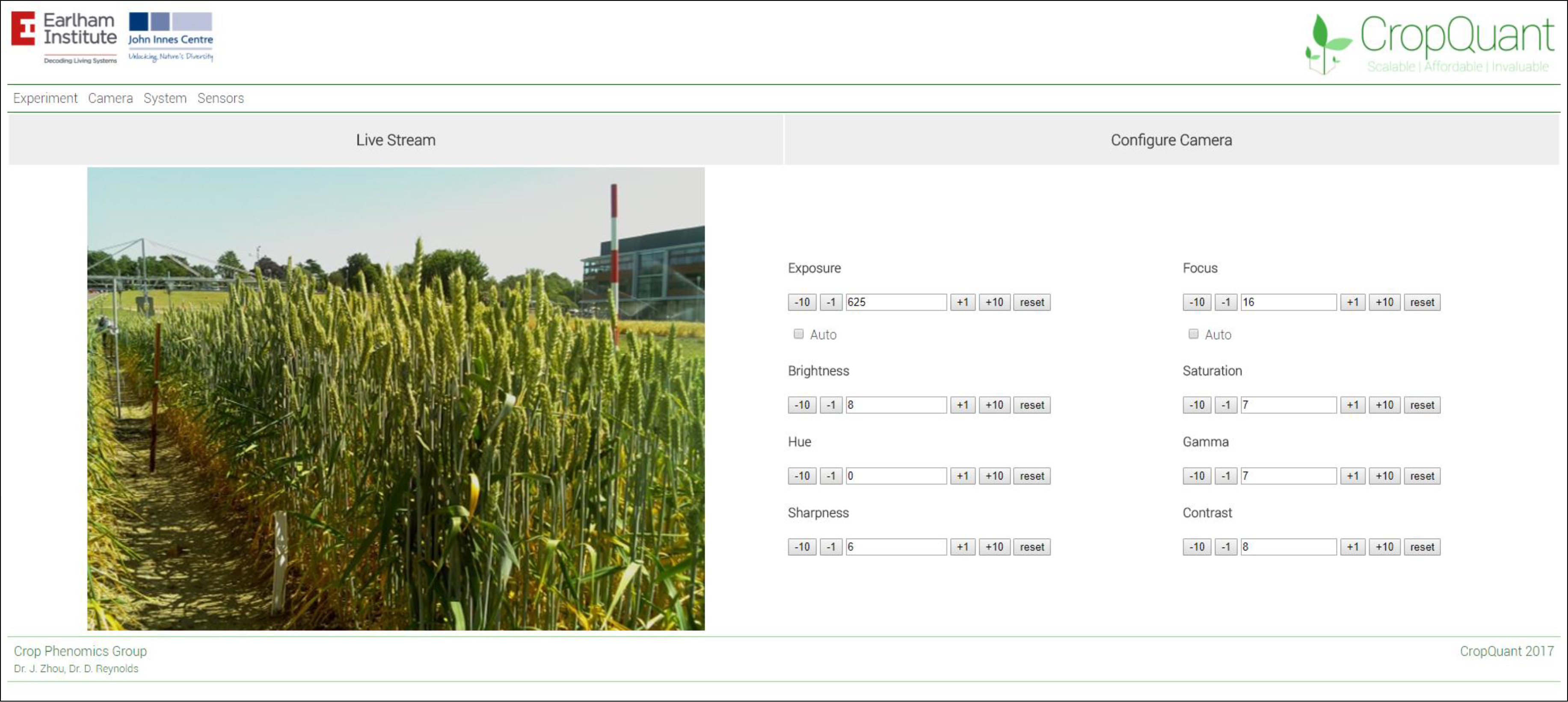
CropSurveyor gives access to each phenotyping device’s interface allowing for device management and configuration such as live video streaming to assist in calibration and experiment setup.

**Supplementary Figure 2:**
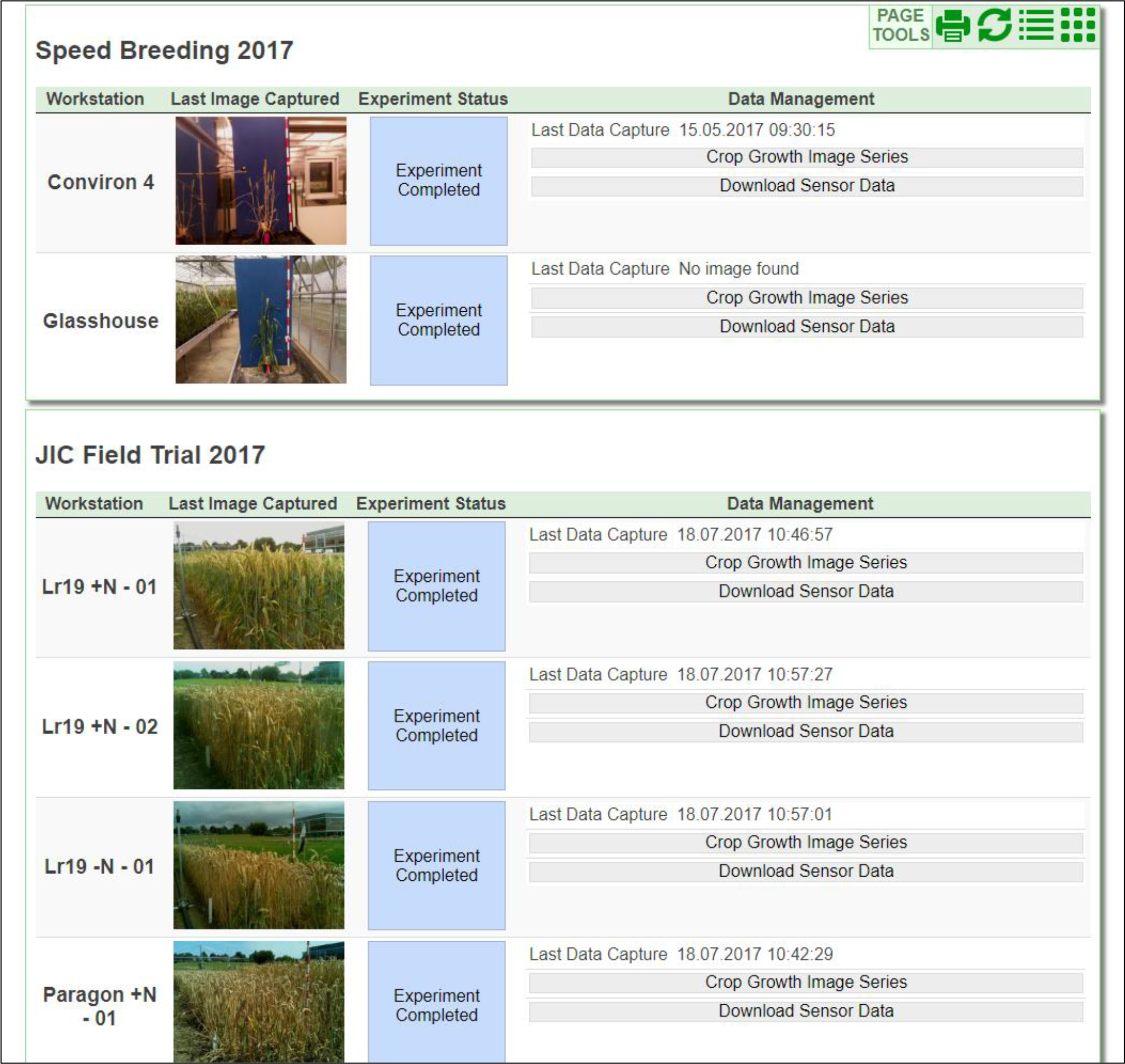
Archived data access allowing browsing and downloading of previous completed trials. Accessing multiple experiments and archived historical data allows cross-referencing data and environmental conditions.

**Supplementary Figure 3:**
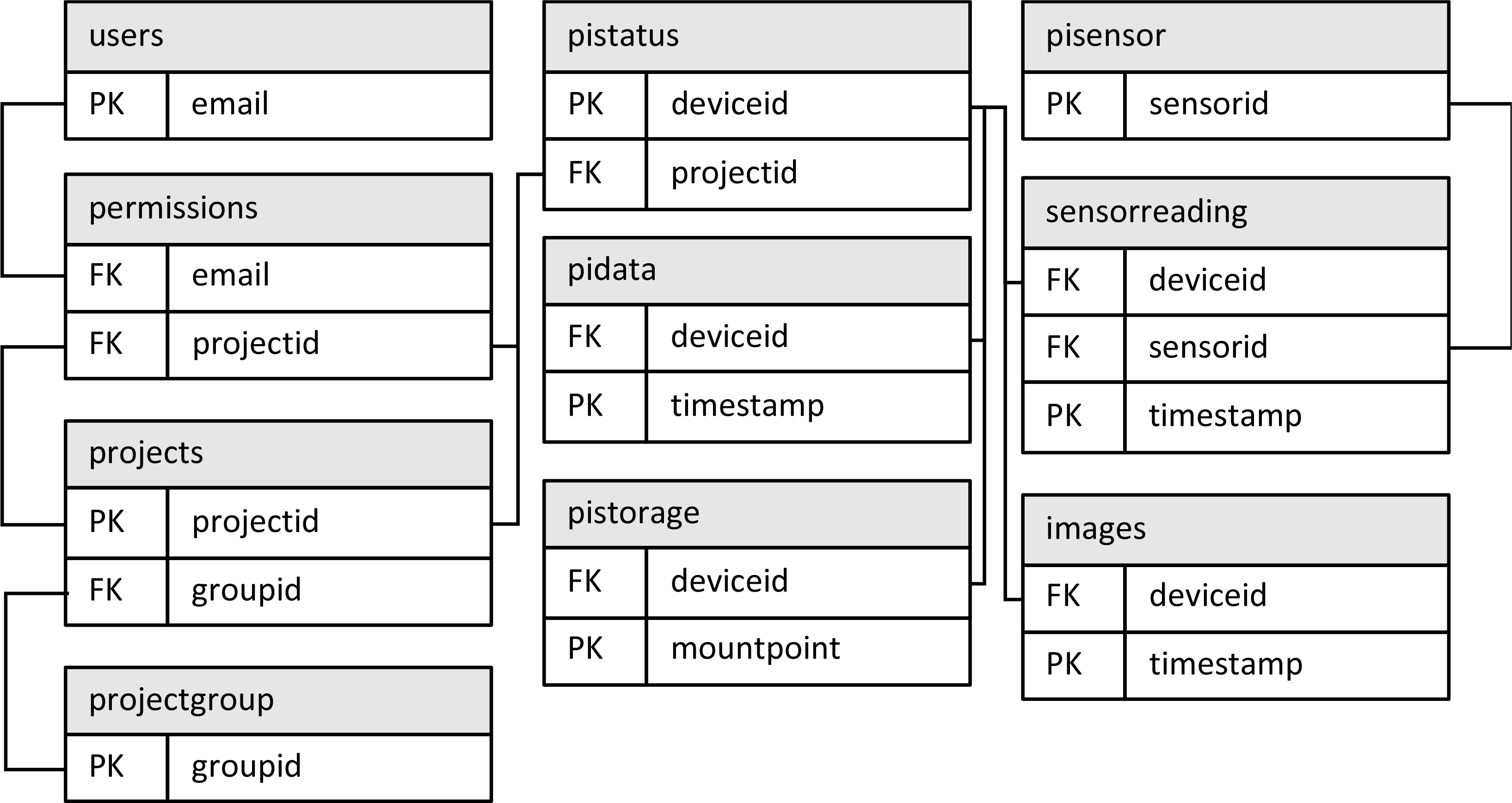
Database Entity-Relationship diagram detailing high level entities within the CropSurveyor database and the relational links between primary, composite and foreign key fields. Diagram describes the structure of database tables; simple storage fields are not shown.

**Supplementary Figure 4:**
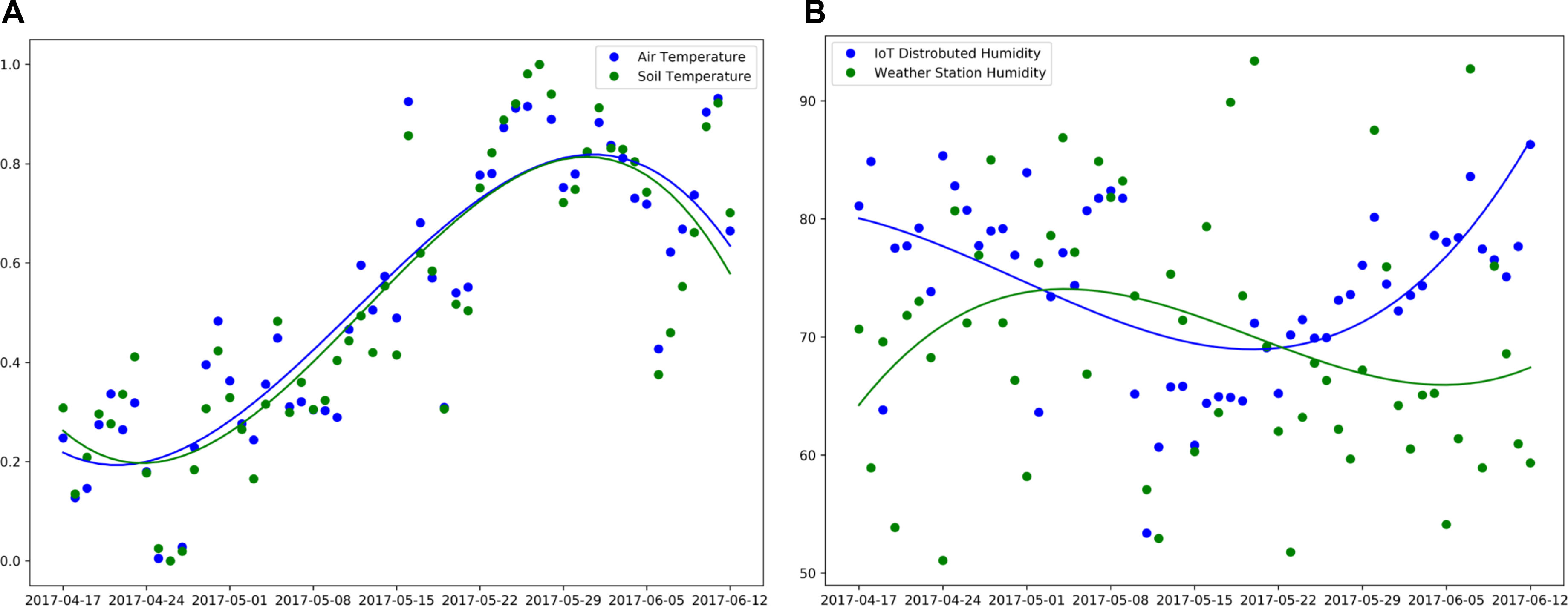
(A) The cross-validation of two different sets of sensors, normalized soil and ambient temperature readings. (B) Different reading between distributed ambient humidity sensors (15 in the field) in comparison with a central weather station, showing different microclimate readings.

## References

1. Tester M, Langridge P. Breeding Technologies to Increase Crop Production in a Changing World. Science (80‐) [Internet]. 2010;327:818–22. Available from: http://www.sciencemag.org/content/327/5967/818%5Cnhttp://www.ncbi.nlm.nih.gov/pubmed/20150489%5Cnhttp://www.sciencemag.org/content/327/5967/818.full.pdf

2. Bevan MW, Uauy C, Wulff BBH, Zhou J, Krasileva K, Clark MD. Genomic innovation for crop improvement. Nature. Nature Publishing Group, a division of Macmillan Publishers Limited. All Rights Reserved.; 2017;543:346–54.

3. Ribaut J-M, de Vicente MC, Delannay X. Molecular breeding in developing countries: challenges and perspectives. Curr Opin Plant Biol. Elsevier Ltd; 2010;13:213–8.

4. Yin X, Struik PPC. Modelling the crop: from system dynamics to systems biology. J Exp Bot. 2010;61:2171–2183.

5. Nagano AJ, Sato Y, Mihara M, Antonio BA, Motoyama R, Itoh H, et al. Deciphering and prediction of transcriptome dynamics under fluctuating field conditions. Cell. Elsevier Inc.; 2012;151:1358–69.

6. Cooper M, Gho C, Leafgren R, Tang T, Messina C. Breeding drought-tolerant maize hybrids for the US corn-belt: Discovery to product. J Exp Bot. 2014;65:6191–4.

7. Reynolds M, Langridge P. Physiological breeding. Curr Opin Plant Biol. Elsevier Ltd; 2016;31:162–71.

8. Reynolds D, Baret F, Welcker C, Bostrom A, Ball J, Cellini F, et al. What is cost-efficient phenotyping? Optimizing costs for different scenarios. Plant Sci. 2018;In Press.

9. Fiorani F, Schurr U. Future scenarios for plant phenotyping. Annu Rev Plant Biol. 2013;64:267–91.

10. Tardieu F, Cabrera-Bosquet L, Pridmore T, Bennett M. Plant Phenomics, From Sensors to Knowledge. Curr Biol. 2017;27:R770–83.

11. Gubbi J, Buyya R, Marusic S, Palaniswami M. Internet of Things (IoT): A Vision, Architectural Elements, and Future Directions. Futur Gener Comput Syst. 2013;29:1645–60.

12. Virlet N, Sabermanesh K, Sadeghi-Tehran P, Hawkesford MJ. Field Scanalyzer: An automated robotic field phenotyping platform for detailed crop monitoring. Funct Plant Biol. 2017;44:143–53.

13. Klukas C, Chen D, Pape J-M. Integrated Analysis Platform: An Open-Source Information System for High-Throughput Plant Phenotyping. Plant Physiol [Internet]. 2014;165:506–18. Available from: http://www.plantphysiol.org/cgi/doi/10.1104/pp.113.233932

14. Vadez V, Kholová J, Hummel G, Zhokhavets U, Gupta SK, Hash CT. LeasyScan: A novel concept combining 3D imaging and lysimetry for high-throughput phenotyping of traits controlling plant water budget. J Exp Bot. 2015;66:5581–93.

15. Humplík JF, Lazár D, Fürst T, Husičková A, Hýbl M, Spíchal L. Automated integrative high-throughput phenotyping of plant shoots: a case study of the cold-tolerance of pea (Pisum sativum L.). Plant Methods. 2015;11:1–11.

16. Kuhlgert S, Austic G, Zegarac R, Osei-Bonsu I, Hoh D, Chilvers MI, et al. MultispeQ Beta: a tool for large-scale plant phenotyping connected to the open PhotosynQ network. R Soc open Sci [Internet]. 2016;3:160592. Available from: http://www.ncbi.nlm.nih.gov/pubmed/27853580%5Cnhttp://www.pubmedcentral.nih.gov/articlerender.fcgi?artid=PMC5099005

17. Busemeyer L, Mentrup D, Möller K, Wunder E, Alheit K, Hahn V, et al. Breedvision ‐ A multi-sensor platform for non-destructive field-based phenotyping in plant breeding. Sensors (Switzerland). 2013;13:2830–47.

18. Zato C, Villarrubia G, Sánchez A, Barri I, Rubión E, Fernández A, et al. PANGEA ‐‐ Platform for Automatic coNstruction of orGanizations of intElligent Agents. In: Omatu S, De Paz Santana JF, González SR, Molina JM, Bernardos AM, Rodríguez JMC, editors. Distrib Comput Artif Intell. Berlin, Heidelberg: Springer Berlin Heidelberg; 2012. p. 229–39.

19. Villarrubia G, De Paz JF, De La Iglesia DH, Bajo J. Combining multi-agent systems and wireless sensor networks for monitoring crop irrigation. Sensors (Switzerland). 2017;17.

20. Neveu P, Tireau A, Hilgert N, Vincent N, Mineau-cesari J, Brichet N, et al. Methods Dealing with multisource and multi-scale information in plant phenomics □: the ontology-driven Phenotyping Hybrid Information System. New Phytol. 2018;

21. Zhou J, Reynolds D, Websdale D, Le Cornu T, Gonzalez-Navarro O, Lister C, et al. CropQuant: An automated and scalable field phenotyping platform for crop monitoring and trait measurements to facilitate breeding and digital agriculture. bioRxiv [Internet]. 2017;1–17. Available from: http://www.biorxiv.org/content/early/2017/07/10/161547

22. UK Government Office for Science. The Internet of Things: making the most of the Second Digital Revolution. London, UK; 2014.

23. Lewandowski CM. Flask Web Development. 1st ed. Eff. Br. mindfulness Interv. acute pain Exp. An Exam. Individ. Differ. Sebastopol: O’Reilly; 2015.

24. Ronacher A. Flask Web Development. BSD licensed; 2018.

25. PHP5 [Internet]. [cited 2018 Oct 10]. Available from: http://php.net

26. Oracle and its affiliates. MySQL 8.0 Reference Manual. 2018.

27. Chen X, Ji Z, Fan Y, Zhan Y. Restful API Architecture Based on Laravel Framework. J Phys Conf Ser. 2017;910.

28. Lindström J, Das D, Mathiasen T, Arteaga D, Talagala N. NVM aware MariaDB database system. 2015 IEEE Non-Volatile Mem Syst Appl Symp NVMSA 2015. 2015;

29. Watson A, Ghosh S, Williams MJ, Cuddy WS, Simmonds J, Rey MD, et al. Speed breeding is a powerful tool to accelerate crop research and breeding. Nat Plants [Internet]. Springer US; 2018;4:23–9. Available from: http://dx.doi.org/10.1038/s41477-017-0083-8

30. Krasner GE, Pope ST. A Description of the Model-View-Controller User Interface Paradigm in the Smalltalk-80 System. J object oriented Program. 1988;1:26–49.

31. Lobell DB. The use of satellite data for crop yield gap analysis. F Crop Res. Elsevier B.V.; 2013;143:56–64.

32. Jones HG. Plants and microclimate: a quantitative approach to environmental plant physiology. Third Edit. Cambridge, UK: Cambridge university press; 2013.

33. White JW, Andrade-Sanchez P, Gore M a., Bronson KF, Coffelt T a., Conley MM, et al. Field-based phenomics for plant genetics research. F Crop Res. Elsevier B.V.; 2012;133:101–12.

34. Chenu K, Cooper M, Hammer GL, Mathews KL, Dreccer MF, Chapman SCS. Environment characterization as an aid to wheat improvement: interpreting genotype-environment interactions by modelling water-deficit patterns in north-eastern Australia. J Exp Bot. 2011;62:1743–1755.

35. King GJ. Crop epigenetics and the molecular hardware of genotype × environment interactions. Front Plant Sci. 2015;6:1–19.

36. Batchelar J, Willets D, Mauley R De, Greening J. A UK Strategy for Agricultural Technologies. 2013.

37. Karp A, Beale MH, Beaudoin F, Eastmond PJ, Neal AL, Shield IF, et al. Growing innovations for the bioeconomy. Nat Plants. 2015;1:15193.

38. Cobb JN, DeClerck G, Greenberg A, Clark R, McCouch S. Next-generation phenotyping: Requirements and strategies for enhancing our understanding of genotype-phenotype relationships and its relevance to crop improvement. Theor Appl Genet. 2013;126:867–87.

39. Roy J, Tardieu F, Tixier-Boichard M, Schurr U. European infrastructures for sustainable agriculture. Nat Plants [Internet]. Springer US; 2017;3:756–8. Available from: http://www.nature.com/articles/s41477-017-0027-3

40. The Government Office for Science. The IoT: making the most of the Second Digital Revolution. WordLink. 2014;1–40.

41. Auernhammer H, Hermann A, Auernhammer H. Precision farming ‐ The environmental challenge. Comput Electron Agric. 2001;30:31–43.

42. Bongiovanni R, Lowenberg-deboer J. Precision Agriculture and Sustainability. Precis Agric. 2004;5:359–87.

43. de Fraiture C, Wichelns D. Satisfying future water demands for agriculture. Agric Water Manag. 2010;97:502–11.

44. Elliott J, Deryng D, Müller C, Frieler K, Konzmann M, Gerten D, et al. Constraints and potentials of future irrigation water availability on agricultural production under climate change. Proc Natl Acad Sci. 2014;111:3239–44.

45. Taylor MC, Hardwick N V., Bradshaw NJ, Hall AM. Relative performance of five forecasting schemes for potato late blight (Phytophthora infestans) I. Accuracy of infection warnings and reduction of unnecessary, theoretical, fungicide applications. Crop Prot. 2003;22:275–83.

